# A generalist microbial predator shows innate attraction to more profitable prey, but does not learn from experience

**DOI:** 10.1101/2024.10.17.618972

**Authors:** P. M. Shreenidhi, Rachel I. McCabe, Heng Liang, Margaret I. Steele, David C. Queller, Joan E. Strassmann

## Abstract

How do generalists adjust to variation in prey abundance and profitability and seek out their preferred prey? We investigated this question in the soil protist *Dictyostelium discoideum*, a generalist predator of many species of bacteria. Despite their generalist diet, amoebas proliferate more quickly on some bacteria than on others. We tested amoeba chemoattraction towards 23 bacterial species and found that they are generally more attracted to the more profitable prey bacteria. Naïve amoebas were also preferentially more attracted to an edible mutant rather than the inedible wild type of a soil *Pseudomonas* isolate. These results suggest that *D. discoideum* amoebas have an innate prey preference that is adaptive. We also tested how experience with different prey bacteria affects chemoattraction in amoebas. Given the huge number of bacterial species in soil, learning from experience should be advantageous. However, we found no evidence that experience with prey bacteria affects preference. Our results suggest that generalist amoebas are innately attracted to the more profitable prey bacteria and this innate attraction cannot be overridden by recent experience.

## Introduction

Predation, where one individual kills and consumes another, is a fundamental interaction between organisms that can profoundly shape organismal evolution. One key component of a predator’s fitness is the ability to sense its prey. Specialist predators are often superior at detecting their prey as compared to generalists (Bernays 2001). Since specialist predators feed on few prey species, they need to track fewer features in their environment to quickly and accurately detect their preferred prey. But generalist predators feed on a diversity of prey species. This very abundance of choice may hinder their ability to detect relevant information about specific prey.

Generalist predators, or more broadly, generalist consumers, can overcome this problem either by innate or learned mechanisms. Generalists might possess innate preferences for certain environmental features, such as shape or colour, that can drive foraging decisions. For example, though honey bees, *Apis mellifera,* harvest pollen and nectar from many flower species, they exhibit an innate preference for some colours (Giurfa et al. 1995). Similarly, the generalist hoverfly, *Eristalinus aeneus,* which feeds on plant pollen and nectar is attracted to artificial floral objects that are bright, radially symmetrical, and possess plant odours (Mishra et al. 2025). However, plasticity in behaviour and the ability to learn from experience are also critical components of foraging behaviour in generalists (Bernays 2001; Mery and Burns 2010). Learning can increase the efficiency of decision making by generalists and allow them to track the most profitable or abundant resources. Honey bees, despite their preference for some colours, can quickly be trained to prefer alternative colours if they are associated with greater rewards (Giurfa et al. 1995). Also, many avian predators learn to avoid chemically defended insects based on their warning coloration (Guilford 1990).

Prey preference has been extensively studied in macroscopic predators such as birds or insects that rely on visual and olfactory cues to track their prey (Guilford 1990; Dukas 2008). These preference behaviours are encoded in the nervous system. But the ability to track prey is crucial even to microscopic organisms that lack a nervous system. Soil protists are often generalist predators that feed on diverse prey bacteria (Johnke et al. 2014; Amacker et al. 2022). Protist predation is a major cause of bacterial mortality in the environment (Jousset 2012). Predation by protists can shape bacterial communities, facilitate nutrient cycling and drive the evolution of bacterial defences (Sherr and Sherr 2002; Jousset 2012; Smith et al. 2023). However, protists use fundamentally different methods for prey detection compared to macroscopic predators.

Prey detection can be challenging for soil protists because of the complexity of soil environments. Soil is a heterogenous environment that varies spatially and temporally in the abundance and diversity of prey bacteria. Even a gram of soil contains extremely diverse micro-organisms and may contain anywhere between two thousand and eight million bacterial species (Gans et al. 2005; Schloss and Handelsman 2006). Soil is composed of minute particles of clay, sand, and decayed plant material separated by water and air-filled pores (Vos et al. 2013).The particles and pores lead to a patchy distribution of resources resulting in fragmented bacterial communities (Raynaud and Nunan 2014; Negassa et al. 2015). Protists must navigate this maze-like soil habitat to seek out their preferred prey bacteria.

*Dictyostelium discoideum*, also known as the social amoeba, is a microbial predator well-known for its fascinating lifecycle which includes both unicellular and multicellular stages (Strassmann and Queller 2011). *D. discoideum* in its simplest form persists as unicellular amoebas which are generalist predators of soil bacteria in temperate forest soils. Prey bacteria that have been co-isolated with *D. discoideum* are diverse and range across at least four bacterial phyla: Proteobacteria, Actinobacteria, Firmicutes and Bacteroidetes (Brock et al. 2018). Despite being generalist predators, these amoebas divide faster when feeding on some prey bacteria than on others (Shreenidhi et al. 2024). Under laboratory conditions, they can double in number in three hours on some of the most edible species of bacteria but take as long as ten hours to double on the less edible species. Some species of bacteria are plainly inedible, while some others are pathogenic (Brock et al. 2018; Steele et al. 2023). Bacteria possess an arsenal of defences to fend off protist predation, including secretion of toxins and chemorepellents, modification of surface properties, and biofilm formation (Jousset 2012; Smith et al. 2023). The edibility of a bacterium could therefore be considered a joint phenotype (Queller 2014) of the amoeba’s offense and the bacteria’s defence. Thus, given the variation in bacterial profitability and defensive responses, it would be advantageous if the amoebas could either innately sense the suitability of a bacterium for predation from a distance or modulate their preference based on their prior experience with a prey bacterium.

*D. discoideum* amoebas use chemotaxis to locate their prey bacteria (King and Insall 2009). Amoebas are attracted to bacterially secreted cyclic adenosine monophosphate (cAMP) (Meena and Kimmel 2017; Rashidi and Ostrowski 2019) and folic acid (Pan et al. 1972; Lima et al. 2014). These two chemoattractants are detected using two different G-protein coupled receptors, cAR1 and fAR1 respectively. There may of course be other, unknown chemoattractants. *D. discoideum* amoebas are also known to be more attracted to Gram-negative bacteria than to Gram-positive bacteria (Rashidi and Ostrowski 2019). However, there is wide-variation in edibility among Gram-negative bacteria (Brock et al. 2018; Steele et al. 2023; Shreenidhi et al. 2024).

In addition to innate preferences, we hypothesized that experience with prey bacteria may play an important role in modulating prey preference in amoebas. The question of whether unicellular organisms can learn from experience and change their behaviour through associative conditioning was a highly disputed topic in the 20^th^ century (Gershman et al. 2021), but there is growing evidence that single-celled protists are sometimes capable of associative learning (Carrasco-Pujante et al. 2021). There are multiple reasons why learning would be beneficial for *D. discoideum* as a generalist predator. Innate preferences may fail to recognize rare species or differentiate between closely related prey that differ in edibility. To account for this, amoebas might learn from experience to be more attracted to profitable bacteria and to avoid unprofitable or harmful bacteria. Prey experience might also direct amoebas towards familiar prey because of the cost of prey-switching. Amoebas experience a reduction in population growth rate when switched from one species of prey bacteria to another (Shreenidhi et al. 2024). These costs are likely physiological and related to changing transcriptional responses to match the new prey bacterium. Thus, it is possible that experience with a prey bacterium primes amoebas to be more attracted towards one bacterium compared to others.

In this study, we measured the chemotaxis of *D. discoideum* amoebas towards 23 phylogenetically diverse species of prey bacteria that were previously isolated from close association with *D. discoideum* from soil environments. We then tested if the chemoattraction of amoebas was explained by the edibility of the prey bacteria. At a finer level, we also tested whether amoebas could differentiate between an inedible *Pseudomonas* strain and an edible transposon mutant of the same species. Next, we tested if amoebas displayed any evidence of learning such that they are differentially attracted to a bacterium when they have prior experience of it being profitable prey.

## Methods

### Population growth rate data

To test for correlation between innate attraction and population growth rate of amoebas on different bacteria, we used previously published population growth rate data of *D. discoideum* for 20 of the 23 bacterial species (Shreenidhi et al. 2024). We used similar assay protocols to estimate population growth rate of *D. discoideum* on our other four bacterial species. We estimated the Malthusian parameter **r** over 20 h on plates.

### Innate prey preference experiments

We performed the innate preference experiments on AX4 amoebas that had been cultured axenically for at least 4 days on HL5 medium including glucose (Formedium) with antibiotics (300 µg/ml Streptomycin, 100 µg/ml Carbenicillin), free of any prey bacteria. By culturing the amoebae axenically we avoided any effects of learned preference for recently experienced bacteria. Amoebas were inoculated at a starting concentration between 5 × 10^5^ – 2 × 10^6^ cells/ml in 25 ml of HL5. The flasks were placed in a shaking incubator at 21°C, 100 rpm. The amoebas were transferred into fresh HL5 media every 3-4 days to maintain them at a concentration below 2 × 10^7^ cells/ml. We collected amoebas for the experiment by centrifuging the cultures at 300*g*, 10°C for 3 minutes. We discarded the spent media and washed the amoeba pellet once more in KK2 buffer (2.25 g KH_2_PO_4_ (Sigma-Aldrich) and 0.67 g K_2_HPO_4_ (Fisher Scientific) per litre). The amoeba pellet was suspended in 2 ml KK2 buffer. We estimated amoeba concentration using a haemocytometer and diluted them to a final concentration of 10^7^ cells/ml for the experiment.

The bacteria for the experiments were cultured on SM/5 (2 g glucose (Fisher Scientific), 2 g BactoPeptone (Oxoid), 2 g yeast extract (Oxoid), 0.2 g MgSO_4_ (Fisher Scientific), 1.9 g KH_2_PO_4_ (Sigma-Aldrich), 1 g K_2_HPO_4_ (Fisher Scientific), 15g agar (Sigma-Aldrich) per litre of deionized water) or LB plates (Fisher Scientific), and resuspended to a concentration of OD_600_ 5 in KK2 buffer.

We used assay protocols similar to Rashidi and Ostrowski (2019) for the chemotaxis experiments. The experiments were performed in 6-well plates containing 2 ml of 2% high purity agar in KK2 buffer per well. In the centre of each well, we pipetted a 2µl droplet of the amoeba suspension (10^7^ cells/ml). A 2 µl spot of the bacterial suspension was placed at 0.65 cm on one side of the amoeba spot and a 2 µl droplet of KK2 buffer at the same distance on the other side (Figure 1A). The spots were placed using a template to keep them identically spaced. We measured the chemotaxis of AX4 amoebas towards 23 species of soil bacteria that had been collected in close association with *D. discoideum,* with three technical replicates per bacterium (Table 1). The experiments were performed over four different days. To account for any day-specific effects, we also performed chemotaxis experiments towards *Klebsiella pneumoniae*, the standard laboratory food given to *D. discoideum*, on all four days. We imaged (phase contrast, 10x magnification) the plates at 6 h from set-up using the Leica Thunder Live Cell Microscope with an automated stage.

**Figure 1.**
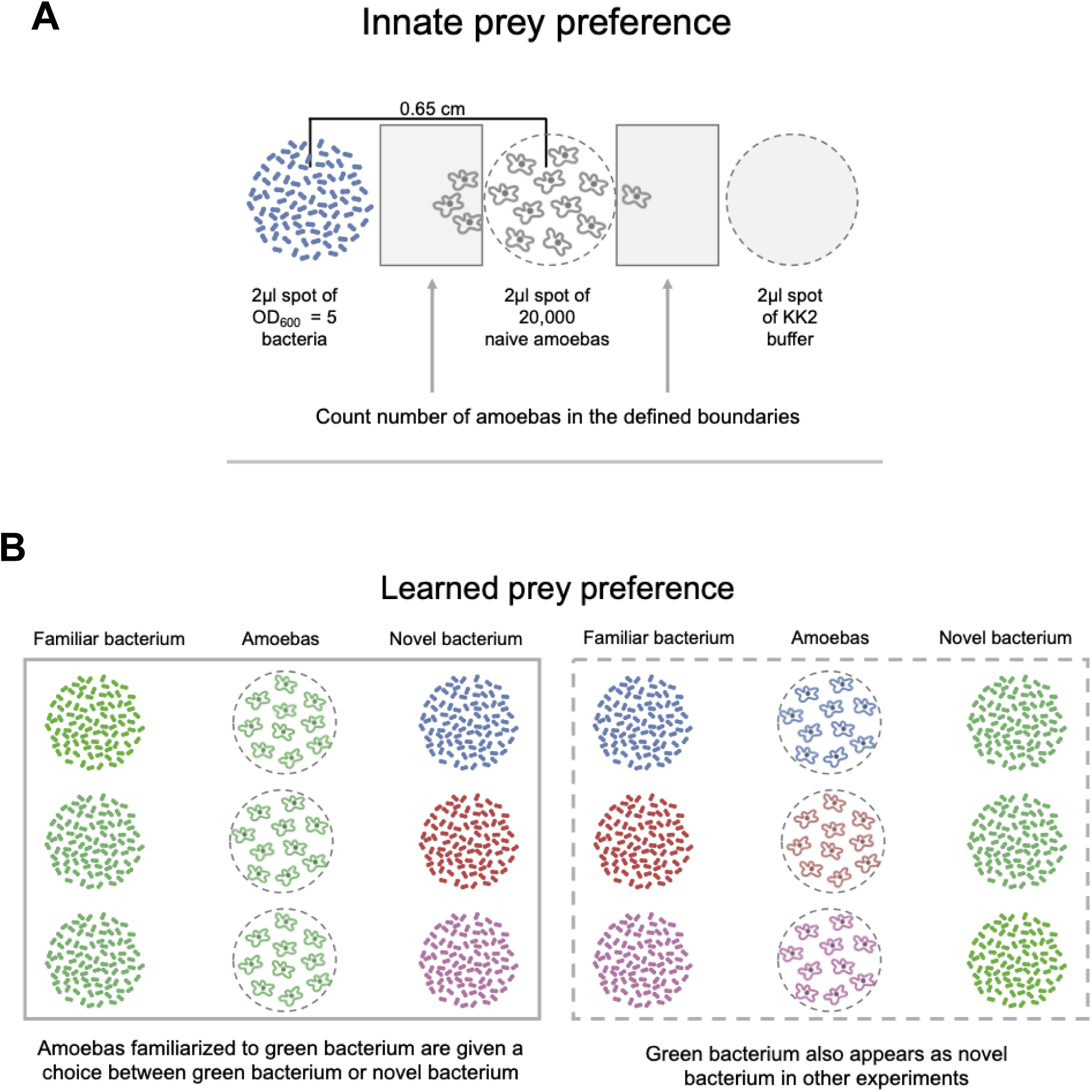
Experimental set-up. A) Innate prey preference in *D. discoideum* amoebas: A spot of AX4 amoebas was placed between spots of bacteria and KK2 buffer. The amoeba spot was imaged after 6 hours to estimate the number of amoebas that move towards the bacteria and buffer. **B) Learned prey preference in *D. discoideum* amoebas**: QS9 amoebas cultured on bacterium A (referred to as familiar bacterium) were given a choice between a familiar and a novel bacterium. The number of amoebas that move towards the familiar and novel bacterium were counted at 4 h and 6 h. Each bacterium appears as both familiar and novel in these experiments. Though we represented only four bacterial species in this figure, the full design included fully factorial tests with eight species.

**Table 1:** List of all bacteria used in innate preference experiments. The bacteria marked in green were also used in learned preference experiments.

| Strain number | Closest species by 16s (Brock2018) | Abbreviation | Phylum/Class | Gram status |
| --- | --- | --- | --- | --- |
| 18P 8.1.1 | *Achromobacter aegrifaciens* | Aa | Betaproteobacteria | Negative |
| 20P 9.1.2 | *Agrobacterium rubi* | Ar | Alphaproteobacteria | Negative |
| 18P 2.2.1 | *Agrobacterium tumefaciens* | At | Alphaproteobacteria | Negative |
| 20P 7.2.1 | *Bacillus aryabhattai* | Ba | Firmicutes | Positive |
| 20P 6.1.2 | *Brucella papionis* | Bp | Alphaproteobacteria | Negative |
| 17D 9.2.1 | *Bacillus toyonensis* | Bt | Firmicutes | Positive |
| 18P 8.2.2 | *Buttiauxella warmboldiae* | Bw | Gammaproteobacteria | Negative |
| 14P 4.3.2 | *Chryseobacterium rhizosphaerae* | Cr | Bacteroidetes | Negative |
| 18P 6.2.3 | *Comamonas testosteroni* | Ct | Betaproteobacteria | Negative |
| 20P 3.2.2 | *Flavobacterium ginsengisoli* | Fg | Bacteroidetes | Negative |
| Lab clone | *Klebsiella pneumoniae* | Kp | Gammaproteobacteria | Negative |
| 11P 7.1.1 | *Mycobacterium arabiense* | Ma | Actinobacteria | Positive |
| 14P 6.2.1 | *Microbacterium maritopicum* | Mm | Actinobacteria | Positive |
| 18P 5.1.3 | *Nocardia coeliaca* | Nc | Actinobacteria | Positive |
| 14P 8.1.1 | *Pseudomonas kuykendallii* | Pk | Gammaproteobacteria | Negative |
| 14P 5.3.2 | *Pseudomonas migulae* | Pm | Gammaproteobacteria | Negative |
| 20P 2.1.2 | *Pandoraea oxalativorans* | Po | Betaproteobacteria | Negative |
| 20P 2.1.1 | *Paenibacillus pabuli* | Pp | Firmicutes | Positive |
| 17P 9.1.1 | *Rhodococcus agglutinans* | Ra | Actinobacteria | Positive |
| 20P 10.2.1 | *Sphingobacterium ginsensidimutans* | Sg | Bacteroidetes | Negative |
| 14P 6.2.3 | *Stenotrophomonas maltophilia* | Sm | Gammaproteobacteria | Negative |
| 5S 2.1.1 | *Staphylococcus saprophyticus* | Ss | Firmicutes | Positive |
| 20P 9.1.1 | *Shinella zoogloeoides* | Sz | Alphaproteobacteria | Negative |
| 20P 10.2.4 | *Variovorax boronicumulans* | Vb | Betaproteobacteria | Negative |

### Innate preference for an edible mutant

We studied the innate attraction of amoebas towards the wild-type *Pseudomonas* strain 20P 3.2.4 (20P324) and its mariner transposon mutant (20P324 6-C11) (Steele et al. 2024). This mutant strain is susceptible to predation by *D. discoideum* amoebas, whereas the wild-type bacterium is inedible (Steele et al. 2024). Using the same assay conditions as described above, we performed experiments where the central spot of AX4 amoebas was flanked on opposite sides by a spot of bacterial suspension (wild type or mutant) and KK2 buffer. We also performed experiments where the wild type and the transposon mutant were paired directly against each other, with the amoeba spot was flanked by both bacterial suspensions. We repeated these experiments on three days with three technical replicates per day. We imaged (phase contrast, 10x magnification) the plates at 6 h from set-up using the Leica Thunder Live Cell Microscope with an automated stage.

### Whole genome sequencing of edible *Pseudomonas* mutant 20P324 6-C11

We extracted Genomic DNA from 20P324 6-C11 cells using the Qiagen DNeasy Blood and Tissue Kit (Qiagen). Illumina sequencing was performed by SeqCenter. Illumina sequencing libraries were prepared using the Illumina DNA Prep kit with a target insert size of 280 bp and sequenced using the Illumina NovaSeq X Plus (2×151bp paired end reads). Bcl-convert (v4.2.4) was used for demultiplexing and adapter trimming. Geneious Prime was used to map reads to the wild type reference genome (NCBI RefSeq assembly GCF_025466075.1), call variants, and exclude variants in low coverage regions. The 20P324 6-C11 genome was assembled using Unicycler (v0.4.9) (Wick et al. 2017), and annotated using the Rapid Annotation using Subsystem Technology (RAST) server (Aziz et al. 2008; Overbeek et al. 2014; Brettin et al. 2015). The mutant was determined to have three mariner transposon insertions not present in the wild type (previously sequenced), which were located in the *gacA* gene, *cyoE* gene, and an in intergenic region between tRNA-dihydrouridine synthase and a gene encoding a hypothetical protein. The edible phenotype is likely caused by the disruption of the *gacA* gene, which positively regulates secondary metabolism (Stallforth et al. 2013).

### Learned prey preference experiment

We studied learned prey preference in a wild strain of *D. discoideum,* QS9, isolated from Mountain Lake Biological Station, Virginia in 2000 and then stored in a -80 °C freezer (Fortunato et al. 2003). To test how experience with a prey bacterium affects prey preference, we first “familiarized” the amoebas to that bacterium. We chose eight bacteria on which *D. discoideum* has a high population growth rate so that the familiar bacterium provides a positive stimulus. To culture amoebas, we spread 200 µl of 1.5 OD_600_ bacterial suspension on SM/5 plates and pipetted 50 µl of *D. discoideum* spore solution containing 200,000 spores.

The amoebas were collected from the resulting vegetative front after 3 - 4 days with a sterile loop. For some bacterial species where the front was hard to see, we collected amoebas from the entire plate after ∼ 36 - 40 hours. We suspended the amoebas in KK2 buffer and centrifuged them at 300*g*, 10 °C for 3 minutes. We washed the amoeba pellet three more times to remove any bacteria. We used the same chemotaxis set-up as described in the “Innate prey preference experiments”, with a central spot of amoebas flanked by two spots of potential attractants. But here the amoebas were given a choice between the bacterium they were cultured on (“familiar bacterium”) and a new bacterium (“novel bacterium”) (Figure 1B). We did this experiment with eight bacterial species in a full factorial design. Amoebas cultured on each of eight familiar bacteria were tested against seven novel bacteria (Table 1, bacterial species highlighted in green). All experiments for amoebas conditioned to a given familiar bacterium were done on the same day, with three technical replicates for every choice between familiar and novel bacterium. We imaged the plates at both 4 h and 6 h from set-up.

### Image Analysis

For all chemotaxis experiments, we used ImageJ version 1.53 (Schneider et al. 2012) to estimate the number of amoebas that moved out of the amoeba spot towards each of the two potential attractants. We drew a line tangential to the amoeba spot at a point that was closest to each potential attractant to define a boundary and counted only those amoebas that moved past this boundary (Figure 1A), as was previously done (Rashidi and Ostrowski 2019). We did the following image processing steps: *Convert to 8-bit > Set Threshold > Make Binary > Fill holes > Apply Watershed*. We manually counted the cells for the innate preference experiments because AX4 amoebas are irregularly shaped and harder to count using automation. We used the *Analyze particles* feature on the relevant parts of the image to count the number of cells for the learned preference experiments that was performed on QS9. Amoebas had not reached the bacteria in any of these experiments, so attraction could not have been caused by feeding amoebas.

## Data analysis & Statistics

### Innate prey preference experiments

We performed all statistical analysis in R (version 4.2.1) (R Core Team 2022). To estimate net amoeba movement towards a given bacterium, we subtracted amoeba movement towards KK2 buffer from amoeba movement towards the bacterial spot. To account for day-specific effects (Figure S1), we scaled all amoeba movement estimated on a given day by dividing it by amoeba movement towards *Klebsiella pneumoniae* (averaged across 3 replicates) on that day. We refer to these values corrected for both buffer and day effects as normalized amoeba movement.

To investigate the relationship between innate preference of amoebas and edibility of the bacterium, we needed to control for the phylogenetic non-independence of data points. We used a phylogeny including all 23 species of soil bacteria used in our experiments as a random factor in our models to control for this. These bacteria are a subset of a larger collection of bacterial strains isolated from close association with *D. discoideum* (Brock et al. 2018). We reconstructed the phylogeny in Brock et al. 2018 by using the previously published 16s sequences of the bacteria (deposited by Brock et al. on GenBank) and 26 species of anchor bacteria (full length 16s sequences downloaded from SILVA rRNA database). A multiple sequence alignment of all sequences was generated using MUSCLE on Geneious Prime. IQ-TREE version 2.3.6 was used to construct a maximum-likelihood tree with model optimization and 100 bootstraps to assess branch support (Minh et al. 2020). We pruned the tree to the 23 species of bacteria used in chemotaxis experiments using iTOL (Letunic and Bork 2021) and used this tree in newick format for all downstream analysis. We forced the phylogenetic tree to be ultrametric using *chronos* function (lambda = 1) in *ape* package in R (Paradis and Schliep 2019) and used *inverseA* function in *MCMCglmm* package (Hadfield 2010) to convert the phylogeny into a covariance matrix.

We then used the R package *MCMCglmm* (Generalized linear-mixed model models using Markov Chain Monte Carlo approach) to examine the relationship between normalized amoeba movement (response variable, henceforth referred to as innate preference) and mean population growth rate of amoebas (fixed effect), controlling for phylogenetic relationships among the bacteria by modelling the bacterial phylogeny as a random effect. We used the default prior for the fixed effect and weakly informative priors for residual variance and random effects (V=1, nu = 0.002) to not overly influence posterior distributions. We report the pMCMC values and the R^2^ for the fixed effect calculated as described using Nakagawa and Schielzeth (2013).

To test for chemokinesis (stimulation of undirected movement), we looked for correlations between average movement towards buffer (response variable) and average movement towards bacterium (fixed effect), scaled as before by average movement towards *Klebsiella pneumoniae* on a given day. We used a MCMCglmm to model this with Bacterial phylogeny as the random factor.

### Innate preference to *Pseudomonas* edibility mutants

To estimate net amoeba movement towards either the wildtype or mutant *Pseudomonas* in buffer experiments, we subtracted amoeba movement towards KK2 buffer from amoeba movement towards the bacterial spot. To test whether net amoeba movement significantly differed based on bacterial identity, we used a Linear Mixed Model (LMM) with bacterial identity (Wild type or Mutant) as a fixed effect, Day as a random effect (Day1, Day2, Day3) and net amoeba movement as the response variable using the R package *lmerTest* (Kuznetsova et al. 2017). To test if amoeba were differentially attracted to either wild-type or mutant bacterium when presented with both, we first calculated proportional amoeba movement to a given bacterium. Next, we tested if arcsine square root transformed proportions were significantly different from 0.5 using a linear mixed model with Day as a random effect.

### Learned preference experiments

We counted the number of amoebas moving towards the familiar bacterium or novel bacterium in each replicate at 4 h and 6 h from set-up. We used the average from these two time points for each replicate for downstream analysis. Replicates with zero amoeba movement were removed from the analysis (2 reps), as they interfere with calculation of proportions. We define *f_i,j_* as the proportional movement of amoebas towards familiar bacterium i, when paired with novel bacterium j, averaged over all three replicates.

We tested if the overall proportional movement (collection of *f_i,j_*’s) was significantly different from 0.5 using a linear mixed effects model with familiar bacterium identity modelled as random effect. We also tested if proportional movement towards familiar bacterium (collection of *f_i,j_*’s for a given i) was dependent on familiar bacterium identity by using a linear model with familiar bacterium identity as fixed effect and arcsine square root transformed proportional movement data as response variable. We used the *emmeans* package to calculate the estimated marginal means and confidence intervals (Lenth et al.

2022).

For a given bacterium *i*, we have two proportional amoeba movement measures, one when the bacterium is familiar (*F_i_*), and another when the bacterium is novel (*N_i_*). *F_i_*, the estimated proportional movement towards bacterium *i* when familiar, averaged over all seven novel bacteria, is calculated as:

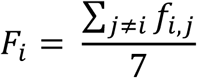

*N_i_*, the estimated proportional movement towards bacterium *i* when novel, averaged over all seven familiar bacteria, is calculated as:

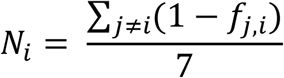

To test if learned preference of amoebas is correlated with innate preference, we built a MCMCglmm model with innate preference towards prey bacterium (data from the innate preference assay) as fixed effect and average proportional movement towards bacterium when familiar (*F_i_*) as the response variable.

To account for innate attractiveness of bacteria when testing if experience with a prey affects chemoattraction of amoebas, we calculated *F_i_* (average movement when familiar) - *N_i_* (average movement when novel) for all 8 species of bacteria and tested if this was significantly different from zero using MCMCglmm. Since these data are also collected across different species of bacteria, we added in the phylogeny of the 8 bacterial species assayed as a random effect. We followed the same steps as described previously to construct the phylogeny, except we pruned the phylogenetic tree down to the 8 species of bacteria used in the learned preference experiments. We used the default priors for the fixed effect and weakly informative prior for residual variance and random effects (V=1, nu = 0.002) to not overly influence posterior distributions. We report the pMCMC values and the R^2^ for the fixed effect calculated as described using Nakagawa & Schielzeth (2013).

## Results

### Amoebas preferentially detect and direct movement towards better prey

To estimate innate prey preference in *D. discoideum* amoebas, we measured the movement of naïve amoebas towards 23 species of soil bacteria or towards KK2 buffer (Figure 1A, Table 1). Amoeba movement was generally directed towards the bacteria and rarely towards buffer (Figure S2). As our measure of innate preference, we subtracted movement towards KK2 from movement towards bacteria and scaled all amoeba movement by movement towards *Klebsiella penumoniae* on the same day to get normalized net amoeba movement towards different species of bacteria (Figure S3). To test if innate response was adaptive, we tested the relationship between normalized movement and population growth rate of amoebas on these bacteria, correcting for the bacterial phylogeny. We found that normalized movement increases with increasing population growth rate (Figure 2; MCMCglmm, Fixed effect posterior mean = 4.58, 95% CI (1.00, 8.11), pMCMC = 0.011, fixed effect R^2^ = 0.21). Thus, amoebas possess an adaptive innate response where they tend to be innately attracted to more profitable bacteria. We found no evidence that this effect dependent on Gram status of the bacteria (Figure S4).

**Figure 2.**
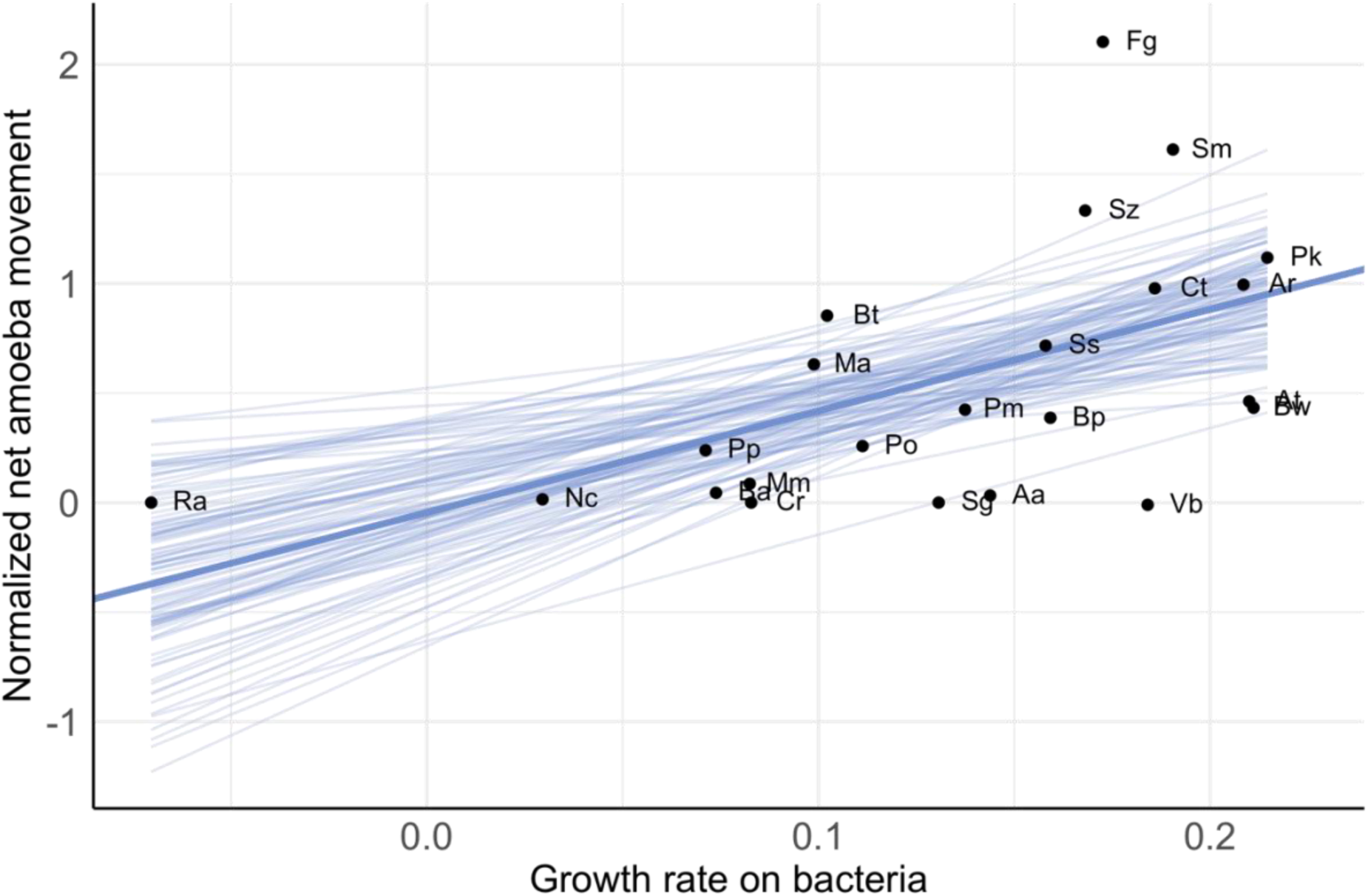
Amoebas are innately attracted to more profitable prey bacteria. Normalized movement of naïve amoebas towards different species of bacteria increases with increasing population growth rate of amoebas on these bacteria (MCMCglmm, Fixed effect posterior mean = 4.58, 95% CI (1.00, 8.11), pMCMC = 0.011, R^2^ = 0.21). Each point is an average of three biological replicates of amoeba population growth rate and three technical replicates of amoeba movement on 23 species of soil bacteria. The solid blue line represents the mean value of slope and intercept from the MCMC glmm model. The fine blue lines are a sampling of 100 slopes and intercepts that lie within the 95% CI distribution of these values from the model.

While chemotaxis is the directed movement of amoebas in response to a chemical gradient, chemokinesis refers to random undirected movement in response to chemical cues. If some bacteria release chemokines that increase amoeba movement overall, then we might observe an increase in amoeba movement in all directions (though presumably weaker at the far end of the amoeba spot). To test if chemokinesis can explain the observed movement patterns, we tested if movement towards KK2 buffer was correlated with movement towards bacteria. We found no significant relationship between these two measures suggesting that these patterns are not driven by chemokinesis (Figure S5: MCMCglmm, Fixed effect posterior mean = 0.06, 95% CI (-0.01, 0.13), pMCMC = 0.12).

### Naïve amoebas are more attracted to an edible mutant over the inedible wild type of a *Pseudomonas* strain

To further test the idea that innate preference of amoebas correlates with bacterial edibility, we measured the chemotaxis of axenically cultured AX4 amoebas towards an inedible *Pseudomonas* soil isolate (20P 3.2.4) and its edible transposon mutant strain (20P324 6-C11). We found that net amoeba movement towards the edible mutant was greater than net amoeba movement (corrected for movement towards buffer) towards the inedible wild type (Figure 3A: Estimate LMM: Estimated net movement (Mutant – WT) = 16.1), t_14_ = 2.38, p-value = 0.03). When amoebas were presented with both bacterial strains simultaneously, we found that the proportion of amoebas that moved towards the edible mutant was significantly greater than 0.5 (Figure 3B: Estimated proportional movement = 0.79, 95 % CI (0.53,0.94), t_8_ = 3.373, p-value = 0.009).

**Figure 3.**
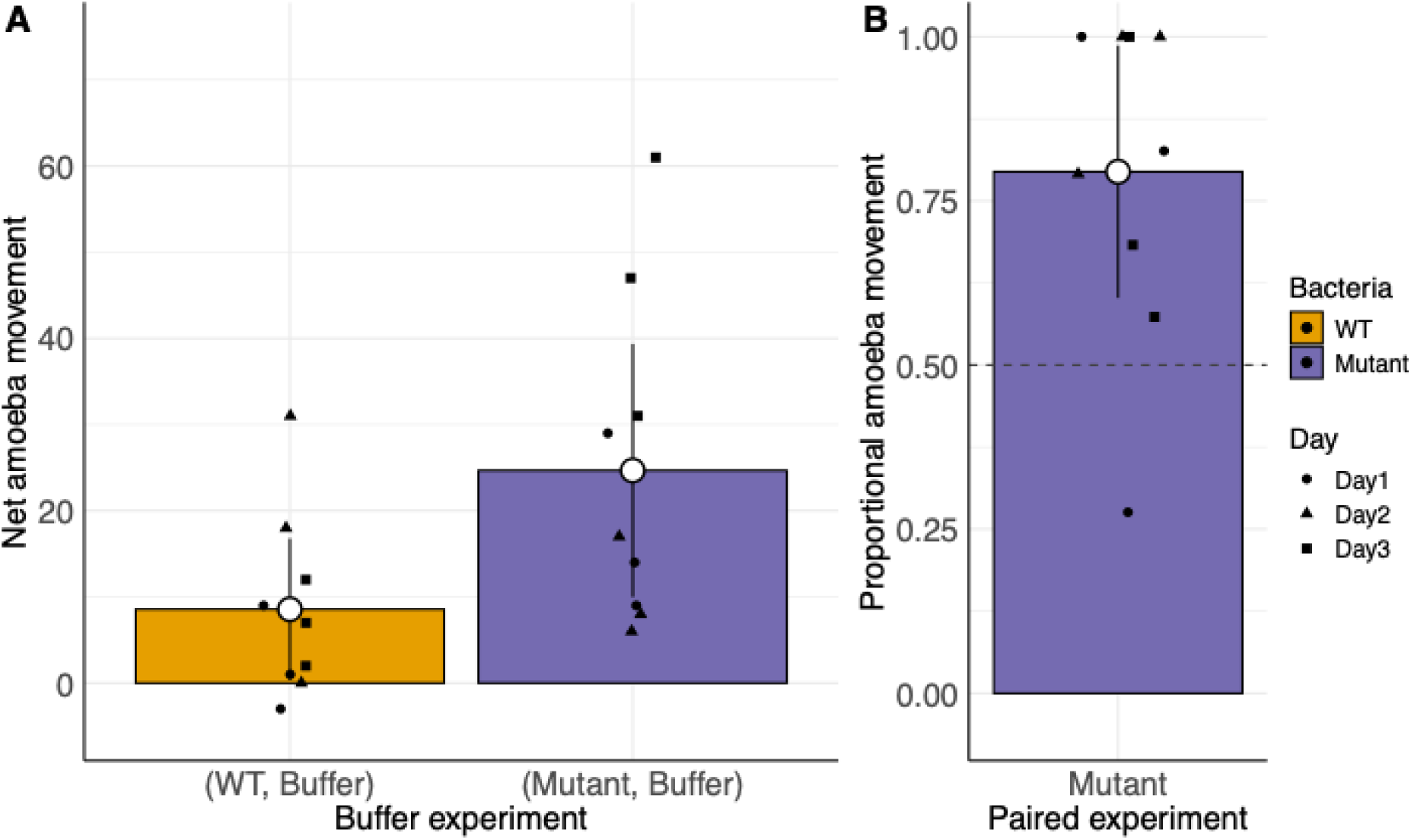
Amoebas prefer the edible transposon mutant over the inedible wild type of *Pseudomonas* sp. 20P 3.2.4. A). In experiments with a choice between a bacterium and **buffer**, net amoeba movement towards the edible mutant is significantly higher than movement towards the wild type *Pseudomonas* bacterium (LMM: p = 0.03). B) In **experiments with a choice between the two bacterial strains**, proportional amoeba movement towards the edible mutant is significantly different from 0.5 (p = 0.009).

### Disruption of the *gacA* gene might be the cause of differential attraction of the amoebas towards *Pseudomonas* bacteria

Whole genome sequencing of *Pseudomonas* 20P 3.2.4 and the mutant revealed transposon insertions in two genes, *gacA* and *cyoE*, and in an intergenic region. The *cyoE* gene is involved in aerobic respiration, while the *gacA* gene is a member of the two-component signal transduction system, GacA/GacS. GacA regulates the transcription of hundreds of genes, including genes involved in the production secondary metabolites that are important for defence against bacterial competitors, plant pathogens and protist predators (Jousset et al. 2009; Hassan et al. 2010; Stallforth et al. 2013). The disruption of the *gacA* gene likely affects the production of many secondary metabolites and is likely to be the cause of increased attraction of amoebas towards the mutant *Pseudomonas*.

### Amoebas do not show a preference for familiar prey

Because the innate effect explains 21% of the variance, there is plenty of scope for improvement via learned effects. To test if prior experience with prey bacteria affects preference in amoebas, amoebas were familiarized to different bacteria and then given a choice between the bacterium they were cultured on (familiar bacterium) or a new bacterium (novel bacterium) (Figure 1B). We chose eight bacteria on which *D. discoideum* has a high population growth rate so that the familiar bacterium provides a positive stimulus, and tested whether amoebas would learn and be more attracted to this good prey. There was no overall trend of preference for the familiar bacterium (Fig 4A: p = 0.476).

**Figure 4.**
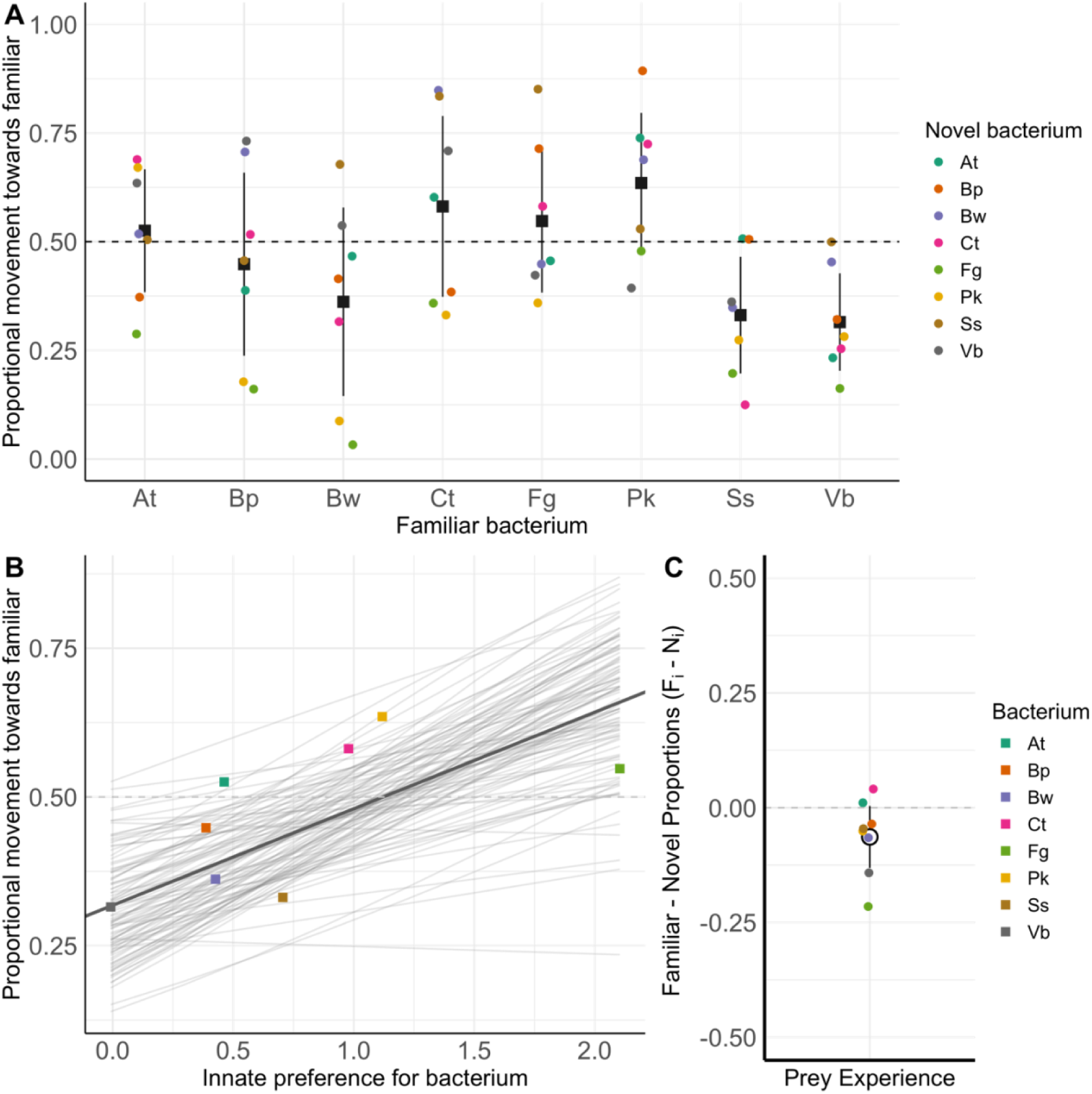
A) Amoebas varied in the level of attraction towards the familiar bacterium. Proportional movement of amoebas towards familiar bacterium averaged across 7 species of novel bacteria. The points represent *f_i,j_* (movement towards familiar bacterium i when paired against novel bacterium j). For each familiar bacterium, the black squares and error bars mark the mean *f_i,j_* across all seven novel bacteria (= *F_i_*) and its 95% confidence interval. **B) The variation in preference for the familiar bacterium was positively correlated with the innate preference for the bacterium** (MCMCglmm, Fixed effect posterior mean = 0.16, 95% CI (-0.02, 0.36), pMCMC = 0.07, R^2^ = 0.28). Here the colored squares represent *F_i_*. **C) Prey experience does not modulate prey preference in *D. discoideum* amoebas.** Difference in proportional movement of amoebas when they are familiar compared to when they are novel (*F_i_ – N_j_*), averaged across all partner bacterial species. There was no significant difference in average movement of amoebas when bacteria are familiar compared to when bacteria are novel. The white circle with the error bars marks the mean plus 95% confidence intervals.

However, we did find wide variation in proportional attraction of amoebas towards the bacterium that depended on bacterial identity (Figure 4A: Familiar bacterium identity: F_8,48_ = 2.81, p-value = 0.012). For example, when amoebas were familiarized to *Pseudomonas sp.* 14P 8.1.1(*Pk*), they seemingly prefer the familiar bacterium (*Pk*) over the novel bacterium (Estimated marginal means = 0.13, 95% CI (-0.006, 0.27), p = 0.06). However, when amoebas were familiarized to *Variovorax sp.* 20P 10.2.4 (*Vb*), they preferred the novel bacterium on average over the familiar bacterium (*Vb*) (Estimated marginal means = -0.18, 95% CI (-0.32, -0.04), p = 0.01).

We found that this pattern of variation in attraction towards the familiar bacterium may be explained by the innate attraction of the amoebas towards the bacterium (Figure 4B; MCMCglmm, Fixed effect posterior mean = 0.16, 95% CI (-0.02, 0.36), pMCMC = 0.07, R^2^ = 0.28). The pattern is not quite significant but, to be conservative, we tested if removing the estimated effects of innate preference affects our results. Because our experiments employ a fully factorial design, each prey bacterium appears in both familiar and novel treatments. Therefore, we accounted for innate preference for the bacterium by comparing the average movement towards bacterium when it is familiar to average movement towards the bacterium when it is novel (*F_i –_ N_j_*) (because innate attraction would be part of both *F_i_* and *N_j_* and cancel out). The difference between the average familiar and novel proportions was not different from zero and the trend is in the direction opposite to learning (Figure 4C; MCMCglmm, Fixed effect posterior mean = -0.08, 95% CI (-0.19, 0.04), pMCMC = 0.151).

## Discussion

All predators must find prey, but generalists that have many suitable prey may have the most difficulty evolving rules to identify them and distinguish degrees of suitability. This may be especially true for microbial predators that encounter many potential bacterial prey species and lack nervous systems to process the information. In this paper, we tested for both innate and learned prey preferences of *D. discoideum* amoebas towards their prey bacteria, as measured by chemoattraction. We found that naïve amoebas are more attracted, on average, to prey bacteria that are more profitable. Thus, innate prey preferences of *D. discoideum* amoebas are generally adaptive.

We also found that such innate preferences can depend on the presence or loss of a single gene in the bacterium. Naive amoebas were more attracted to an edible mutant of the 20P 3.2.4 *Pseudomonas* strain compared to the inedible wild type. The mutant differed at three sites but the likely causal mutation was a transposon insertion in the *gacA* gene. This gene positively regulates secondary metabolism, and is important for predation resistance in many species of *Pseudomonas* including the *Pseudomonas* strain 20P 3.2.4 (Stallforth et al. 2013; Inglis et al. 2018; Steele et al. 2024). 20P 3.2.4 is predicted to have 13 secondary metabolite biosynthetic gene clusters based on analysis with antiSMASH (Steele et al., 2023). Targeted deletion of the *gacA* gene in 20P 3.2.4 is known to increase susceptibility to predation by amoebas similar to the increase seen in the transposon mutant (Steele et al. 2024). It is possible that secondary metabolites that confer predation resistance also act as chemorepellents. Thus, bacteria may be signalling their unsuitability for predation using some secondary metabolites much like the way many insects signal their unsuitability as prey using warning colouration. It is also possible that *gacA* downregulates production of the normal chemoattractants. Either way, the result suggests modulating chemoattraction is a novel function of *gacA*.

The positive relationship between innate preference and prey profitability hints at the long evolutionary history of amoeba-bacteria interactions and how these innate responses have been selected to enhance fitness over evolutionary time. But as with any correlation we cannot establish the direction of causation between innate preference and prey profitability. It seems likely that variation in amoeba performance on different bacteria shaped prey preferences. But it is also possible that amoebas improved their performance on bacteria that were already attractive.

Our data shows that only 21 % of variance in amoeba attraction is explained by prey profitability suggesting that there are other constraints on this response. For example, *D. discoideum* amoebas use chemotaxis not only to locate their bacterial prey, but also to locate other conspecific amoebas during starvation to initiate the multicellular cycle. Thus, innate chemoattraction towards bacterial prey cannot be changed to the extent that it hinders the ability of the amoebas to find other conspecifics. Indeed, cAMP is a common chemoattractant signal used by *D. discoideum* to detect both bacterial prey and conspecifics using the chemoreceptor cAR1 (Klein et al. 1988; Meena and Kimmel 2017; Rashidi and Ostrowski 2019). A greater constraint on optimal amoeba preferences is evolution of the prey species. Because prey bacteria are constantly evolving, it seems impossible to evolve innate preferences that remain adaptive on all encountered bacterial species. Prey would presumably be under selection to become “invisible” by reducing secretion of attractants, and might also evolve mimicry of secretions of unattractive bacteria. Thus, whatever the reasons for the imperfection of innate attraction, there would likely be considerable advantage to adjust preferences via learning.

However, we found that prior experience with a prey species did not generally seem to influence subsequent attraction of *D. discoideum* amoebas to that bacterium. Because these were all profitable prey species, we expected amoebas might learn to be more attracted after experiencing them, but they did not. The idea that amoebas can change their prey preference based on experience rests on the assumption that the prey bacteria secrete different chemoattractants, or combinations of chemoattractants, and that amoebas can distinguish among them, probably by using different chemoreceptors or combinations of chemoreceptors. Experience with a prey bacterium could then lead to upregulation of those specific chemoreceptors, or to give greater weight to information from those chemoreceptors, thus reinforcing preference. The only prey chemoattractants of *D. discoideum* that are well understood are folate and cAMP (Artemenko et al. 2014). The response to these chemoattractants is mediated by two highly-ligand specific G-protein coupled receptors (GPCRs): the cAR1 receptor for cAMP and the fAR1(GrlL) receptor for folate (Pan et al. 2016; Meena and Kimmel 2017). But amoebas possess 61 putative receptors in the GPCR family (Heidel et al. 2011). Many of these potential GPCRs are of unknown function and some could be undiscovered chemoreceptors that play a role in modulating responses based on experience. All that would be needed is a GPCR with a different ligand binding domain to activate the signalling machinery that stimulates actin remodelling to trigger chemotaxis.

Though we observed no consistent change in amoeba response based on experience with prey, variation in amoeba movement in learning experiments was partly explained by their innate preference for a specific bacterium. It is possible that the bacteria assayed in our experiments do not secrete sufficiently different chemoattractants, or that amoebas only use folate and cAMP to detect their prey bacteria. We know that amoeba attraction towards these two chemoattractants is dose-dependent where higher doses of the attractant lead to more amoeba movement (Meena and Kimmel 2017; Rashidi and Ostrowski 2019). Thus, it is possible that the prey bacteria we tested vary in the levels of cAMP or folate production and that amoebas simply prefer prey bacteria that produce more cAMP or folate, irrespective of prey experience. However, the lack of observed learning may have explanations other than an absence of multiple chemoattractants.

The levels of environmental heterogeneity experienced by an organism play an important role in the evolution of innate and learned responses (Mery and Burns 2010). Innate behavioural responses evolve when the environment is relatively stable over several generations, while plastic/learned responses evolve when the environment changes rapidly over the organism’s lifetime (Stephens 1991). The presence of an innate adaptive prey preferences and the lack of evidence for learning in our experiments suggests that perhaps amoebas do not experience very much environmental heterogeneity in prey species. This seems surprising given the diversity of bacteria in the soil but the micro-distribution of prey species is worth further investigation.

The importance of self-generated gradients in long-range movement may be a more likely explanation for why we observed no learning. Chemotactic gradients established solely by the diffusion of attractants from a local source are only effective at guiding directional movement of cells over short distances (500 μm) (Tweedy et al. 2016). The gradient might be too shallow to establish any directional movement over large distances. At long distances (5 mm) self-generated gradients play an important role (Tweedy et al. 2016). Self-generated gradients occur when cells produce enzymes that break down the chemoattractant locally. This can steepen the attractant gradient and allow directional movement towards the source. *D. discoideum* amoebas secrete deaminases to break down folate (Bernstein et al. 1981) and cAMP phosphodiesterase to breakdown cAMP (Shaulsky et al. 1998). Our experiments involve distances well above 500 μm, so attractant breakdown is likely to be quite important. Amoeba movement in this context might be limited to chemoattractants that can be degraded such as folate and cAMP. In experiments using much shorter distances, amoebas may be able to respond to, and perhaps learn from, chemoattractants that they cannot degrade.

In conclusion, our study shows that innate preferences are a key component of foraging behaviour in a microbial predator, *Dictyostelium discoideum*. We found no evidence for learned prey attraction behaviours. This is a puzzling result that merits further exploration. Future work could focus on the microstructure of bacterial soil communities experienced by amoebas and if that is conducive for learning. Further characterization of GPCRs could reveal whether or not the amoebas possess a flexible mechanism for in sensing different chemoattractants. Finally, experiments involving attraction over shorter distances might uncover learned foraging behaviours at smaller scales that we tested.

## Author contributions

P.M.S., D.C.Q., and J.E.S. designed research. P.M.S., R.I.M., H.L., and M.I.S., performed research. P.M.S., R.I.M., H.L., M.I.S., D.C.Q., and J.E.S. analyzed the data. P.M.S., D.C.Q., and J.E.S. wrote the paper with inputs from R.I.M., H.L., and M.I.S.

## Acknowledgements

This material is based upon work supported by the National Science Foundation (DEB 1753743 and DEB 2237266). We thank Debra Brock for inputs during the development of the project. Robert Insall provided insight on short versus long-range chemoattraction. We also thank members of the Queller-Strassmann lab for useful discussions.

## Supplementary figures

**Figure S1:**
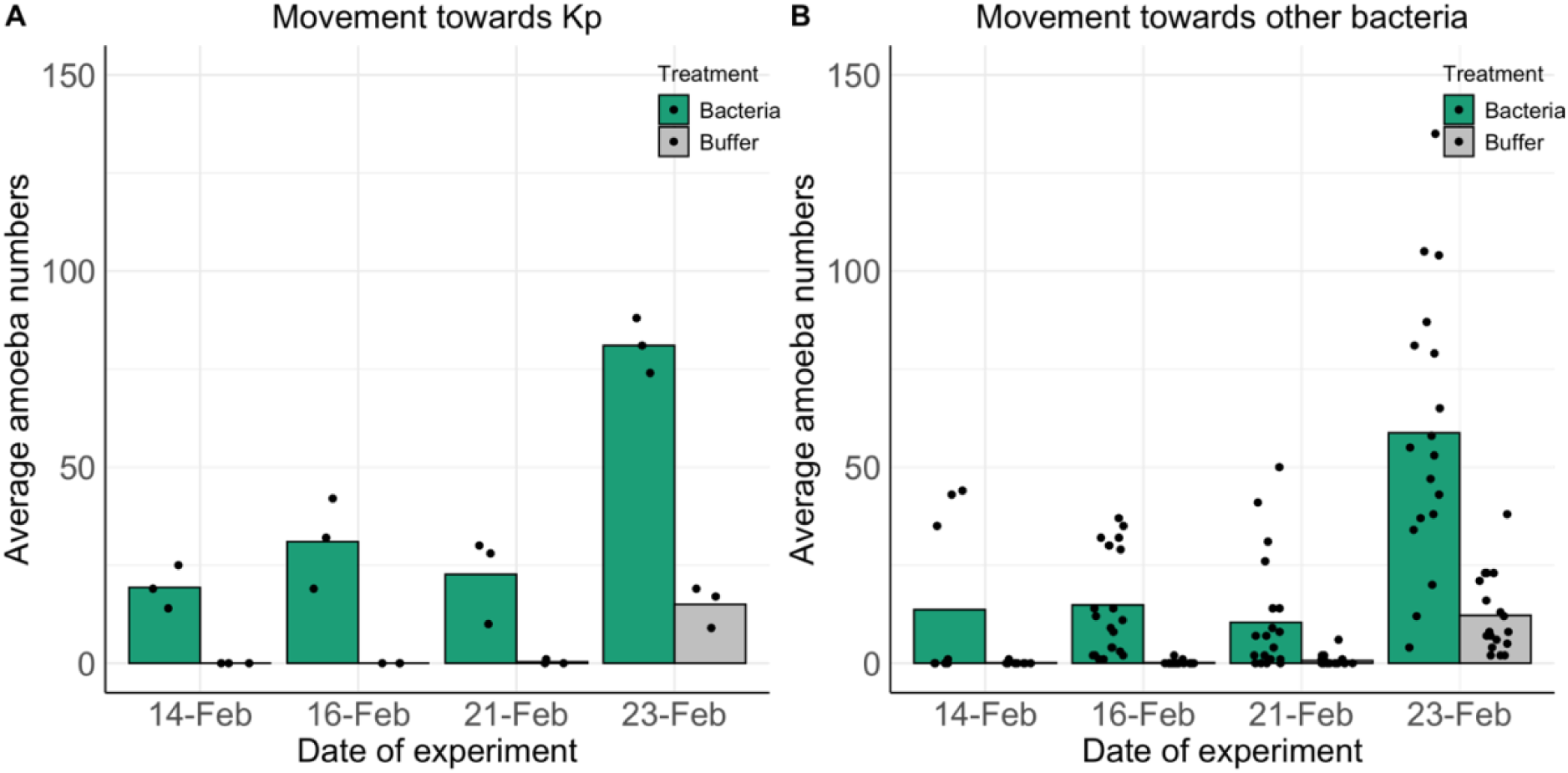
A) Movement of AX4 amoebas towards *Klebsiella penumoniae* on different days. B) Movement towards other bacteria on different days. Since chemotaxis experiments towards *Klebisella penumoniae* were repeated on all days, we can see that there are day-specific effects that need to be corrected for.

**Figure S2:**
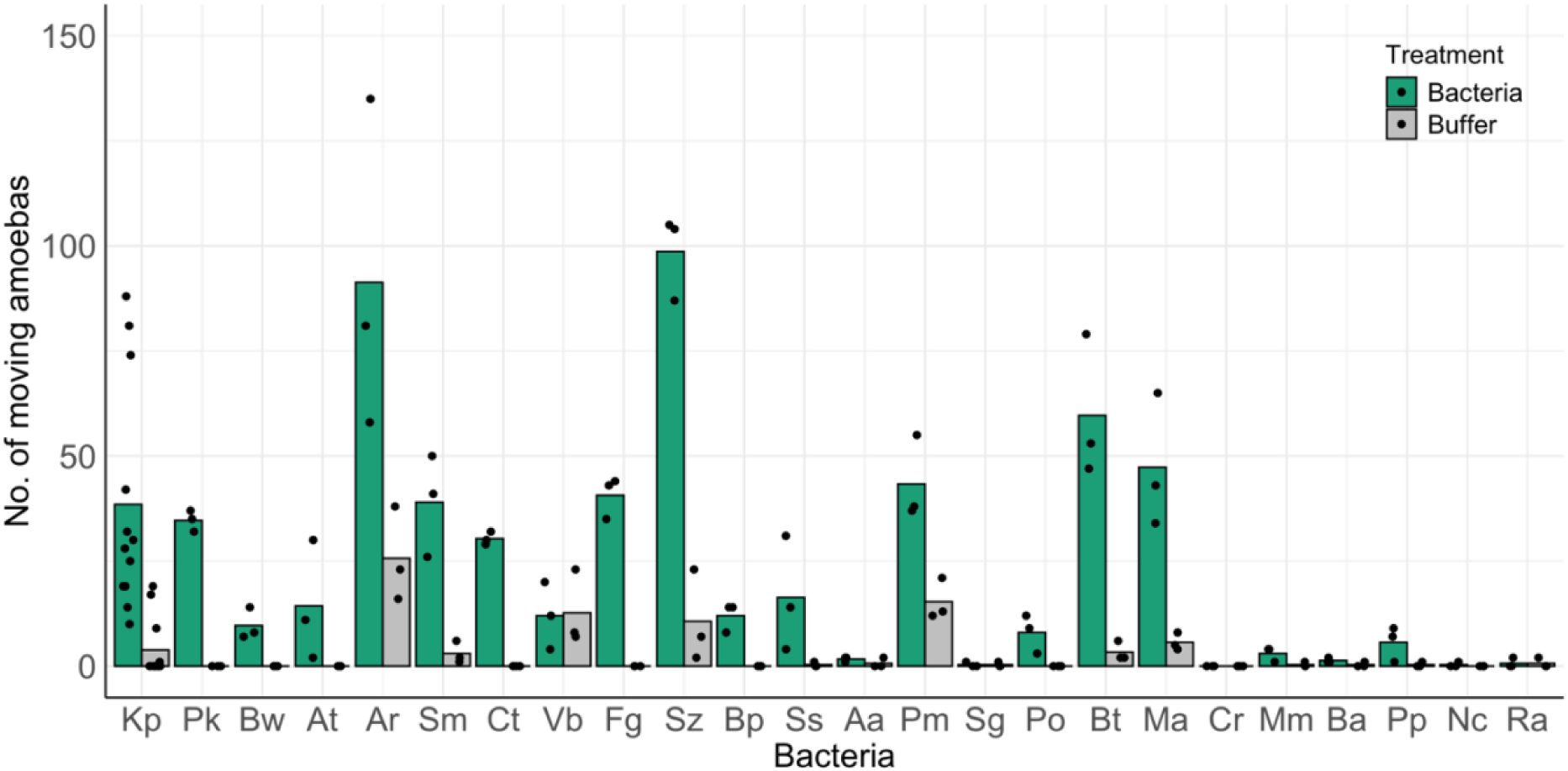
Data from innate preference experiments. The movement towards the bacterial spot is represented by green bars, and movement towards KK2 buffer is represented by grey bars. Amoebas generally move towards the bacteria and rarely to buffer. The bar marks the means across the 3 technical replicates points (more for the *Kp* control).

**Figure S3:**
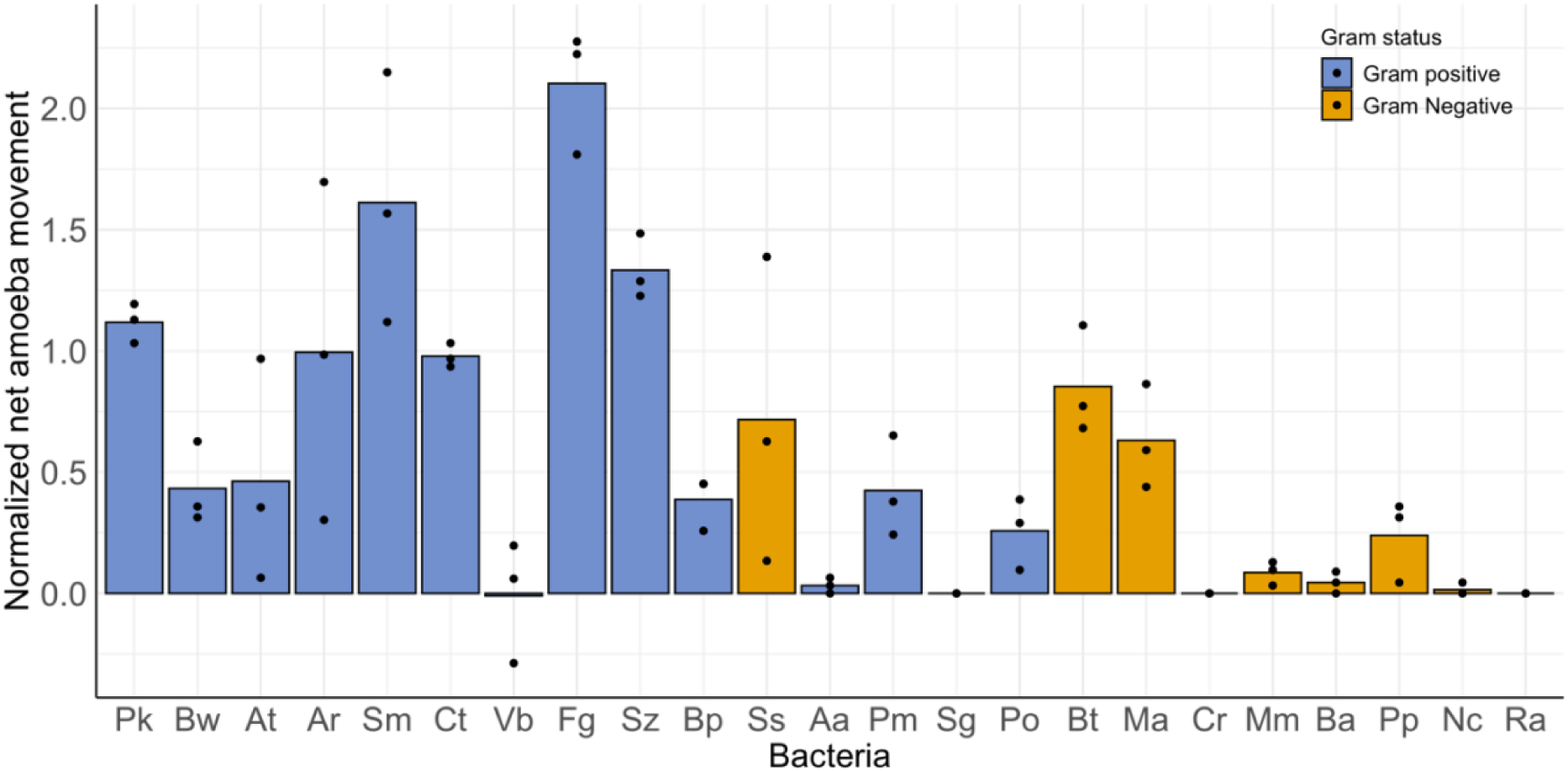
Normalized movement of AX4 amoebas towards different species of bacteria. We first calculated net amoeba movement towards a bacterium in each replicate by subtracting the number of amoebas that moved towards KK2. To make amoeba movement comparable across different days, we divided net movement towards bacterium by average net movement towards *Klebsiella pneumoniae* on that day. Bar marks the mean across 3 technical replicates (points).

**Figure S4:**
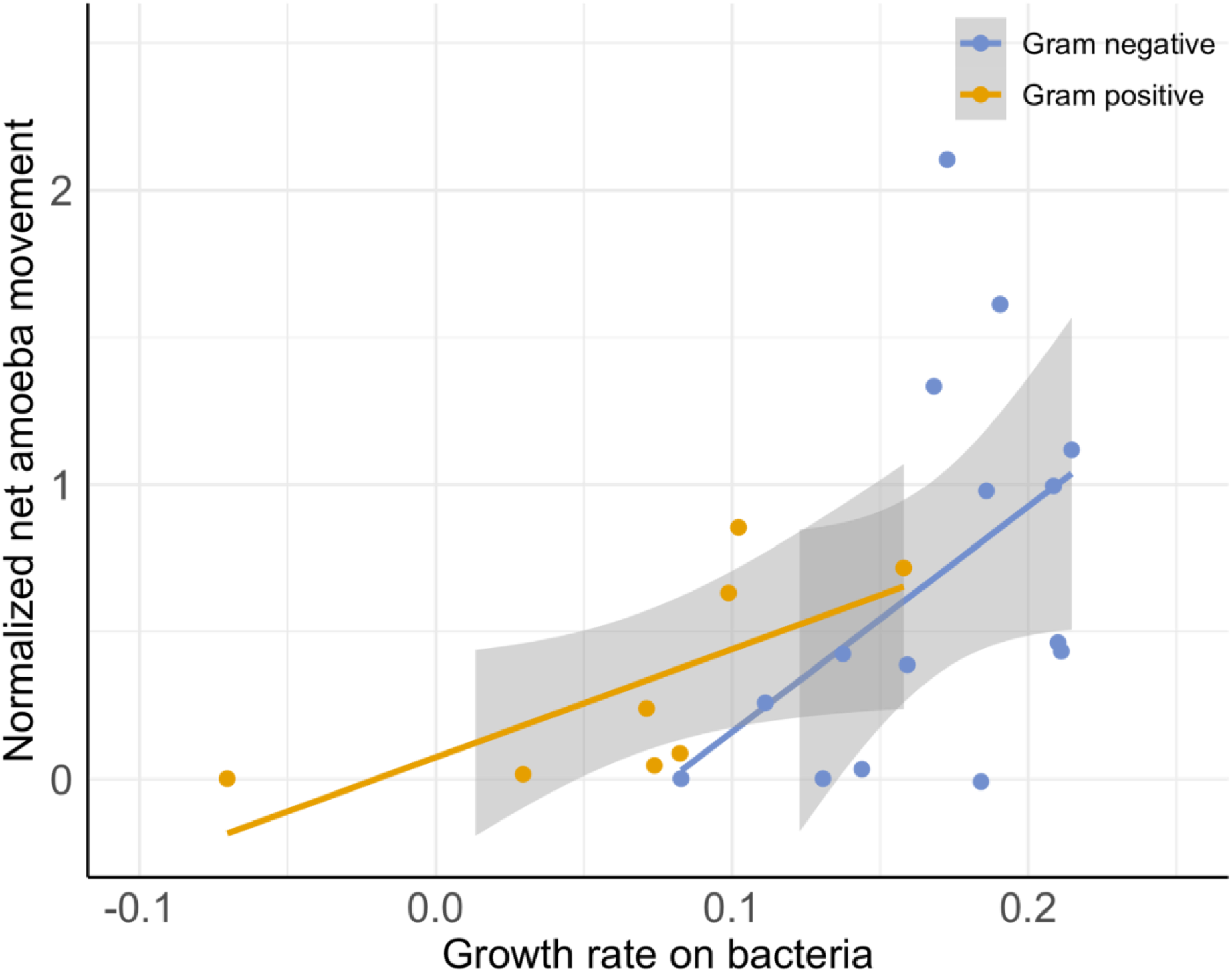
Normalized movement of amoebas versus their growth rate on various bacteria. We found no significant effect interaction effect between Gram status and mean growth rate on normalized amoeba movement (p = 0.37). We used a linear model with normalized amoeba movement as the response variable, and included mean amoeba population growth rate and bacterial Gram status as fixed effects, along with their interaction.

**Figure S5:**
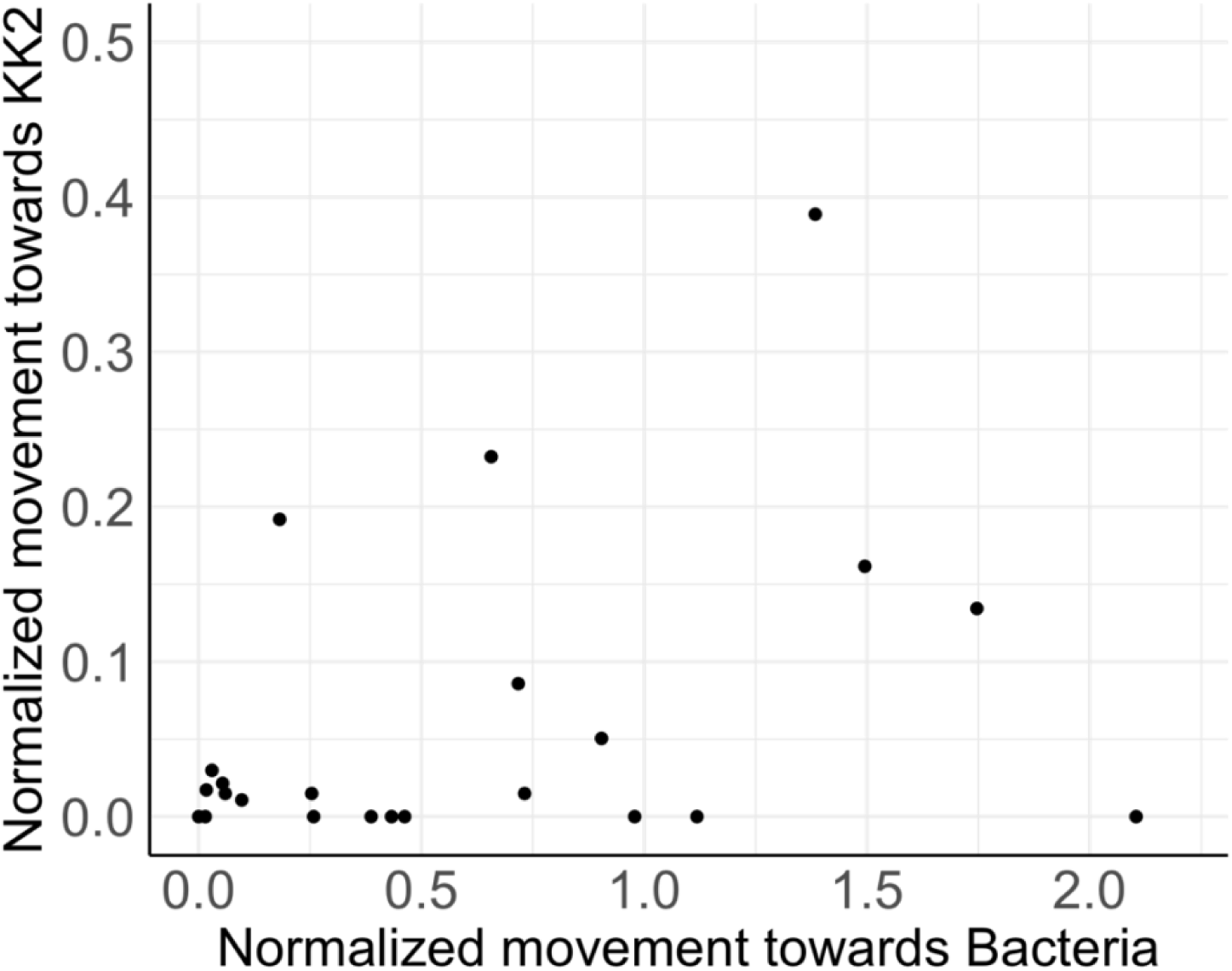
Normalized movement towards KK2 buffer vs. Normalized movement towards bacteria. Here normalization means correction for any day effect, where we divided all movement on a given day by mean movement towards *Klebsiella pneumoniae* on that day. We found no significant relation between movement towards buffer and movement towards bacteria (Fixed effect posterior mean = 0.06, 95% CI (-0.01, 0.13), pMCMC = 0.12).

